# Chiral evasion and stereospecific antifolate resistance in *Staphylococcus aureus*

**DOI:** 10.1101/2020.07.28.224667

**Authors:** Siyu Wang, Stephanie M. Reeve, Adegoke A. Ojewole, Marcel S. Frenkel, Graham T. Holt, Pablo Gainza, Santosh Keshipeddy, Vance G. Fowler, Dennis L. Wright, Bruce R. Donald

## Abstract

Antimicrobial resistance is a health care crisis. The resistance-conferring mutation F98Y in *Staphylococcus aureus* dihydrofolate reductase (SaDHFR) reduces effectiveness of antifolates, e.g., trimethoprim (TMP). Although propargyl-linked antifolates (PLAs) are much more resilient than TMP towards F98Y, this substitution still vitiates their inhibition potency. Surprisingly, differences in the enantiomeric configuration at the stereogenic center of PLAs influence the isomeric state of NADPH cofactor. Is resistance correlated with chiral evasion? A mechanism of action underpinning this coupling is unknown. To understand the molecular basis of F98Y-mediated resistance and how PLAs’ inhibition drives NADPH isomeric states, we used OSPREY to analyze a comprehensive suite of structural, biophysical, biochemical, and computational data. We present a model showing how F98Y SaDHFR exploits a different anomeric configuration of NADPH to evade certain PLAs’ inhibition, while other PLAs remain resilient to resistance. Our model should enable general design of inhibitors that are resilient to chiral evasion.

## 1 Introduction

Methicillin-resistant *Staphylococcus aureus* (MRSA) is one of leading causes of healthcare associated infections, pneumonia and skin and soft tissue infections (SSTIs). Antifolates such as trimethoprim (TMP) exhibit potent activity as antibacterial agents against many MRSA clinical isolates (*1, 2*). Due to structural and sequence differences between human and prokaryotic dihydrofolate reductase (DHFR) and the specific effects on quickly replicating cells, antifolates can be developed as selective and safe antimicrobial agents (*3*). Currently a fixed-dose combination of TMP and sulfamethoxazole, a dihydropteroate synthetase inhibitor, is one of the top ten oral antibiotics prescribed (*4*).

DHFR is an enzyme vital for cellular replication, mainly due to its important role in the one-carbon metabolism pathway (*5*). DHFR reduces dihydrofolate (DHF) to tetrahydrofolate (THF) (*6*). Nicotinamide adenine dinucleotide phosphate (NADPH) is required for this catalytic reaction. NADPH participates as a cofactor and donates a hydride to reduce C4 on the dihydropterin ring of DHF. Over the years, various models for different mechanisms of catalysis have been proposed (*7–9*). During catalysis, NADPH adds a hydride to C6 of the dihydropterin ring and then N5 is protonated (*9, 10*). In addition, in *Escherichia coli* DHFR, Tyr98 stabilizes the positive charge in the nicotinamide ring during the hydride transfer between C4 of the nicotinamide ring and C6 of the dihydropterin ring (*11*). In wild type SaDHFR, position 98 is a phenylalanine. Its mutation to tyrosine confers resistance to antifolates such as TMP and some other novel antifolates according to experimental and clinical observations (*12, 13*).

New resistance mechanisms continue to emerge and challenge the efficiency of existing drugs (*14*). Resistance to antifolates, including TMP, the only approved antifolate for antibiotic use, has emerged in many cases (*12, 15*). Therefore, second generation antifolates that can overcome resistance must be developed. This need motivates our studies to develop and apply rapid and effective computational methods to predict and understand the mechanisms of resistance.

Recently we have developed a series of propargyl-linked antifolates (PLA) that are potent SaDHFR inhibitors (*13,16*). The key feature of the PLA class is its eponymous propargyl linker (see Table 1). As with TMP, PLAs are competitive inhibitors that comptete with DHF to bind in the active site of DHFR. The linear triple bond linker makes it possible for PLAs to fit into the narrow binding pocket with limited steric clashes (see Table 1 and Figure 1a). PLAs also have a biaryl ring system that mimics the para-aminobenzoic acid (PABA) and glutamate moieties of folate. In contrast to traditional antifolates such as TMP, PLAs showed high potency against both wild type and predominant TMP-resistant strain enzymes according to IC_50_ measurement (*13, 16*).

**Table 1:** IC_50_ data of R-27, S-27 and TMP.

| Inhibitor | Sa IC<sub>50</sub> (nM) | Sa(F98Y) IC<sub>50</sub> (nM) |
| --- | --- | --- |
| R-27 | 15 ± 0.7 | 510 ± 50 |
| S-27 | 18 ± 2 | 111 ± 6 |
| TMP | 23 ± 3 | 1700 ± 19 |

Adapted from Table 2 in (16).

**Figure 1:**
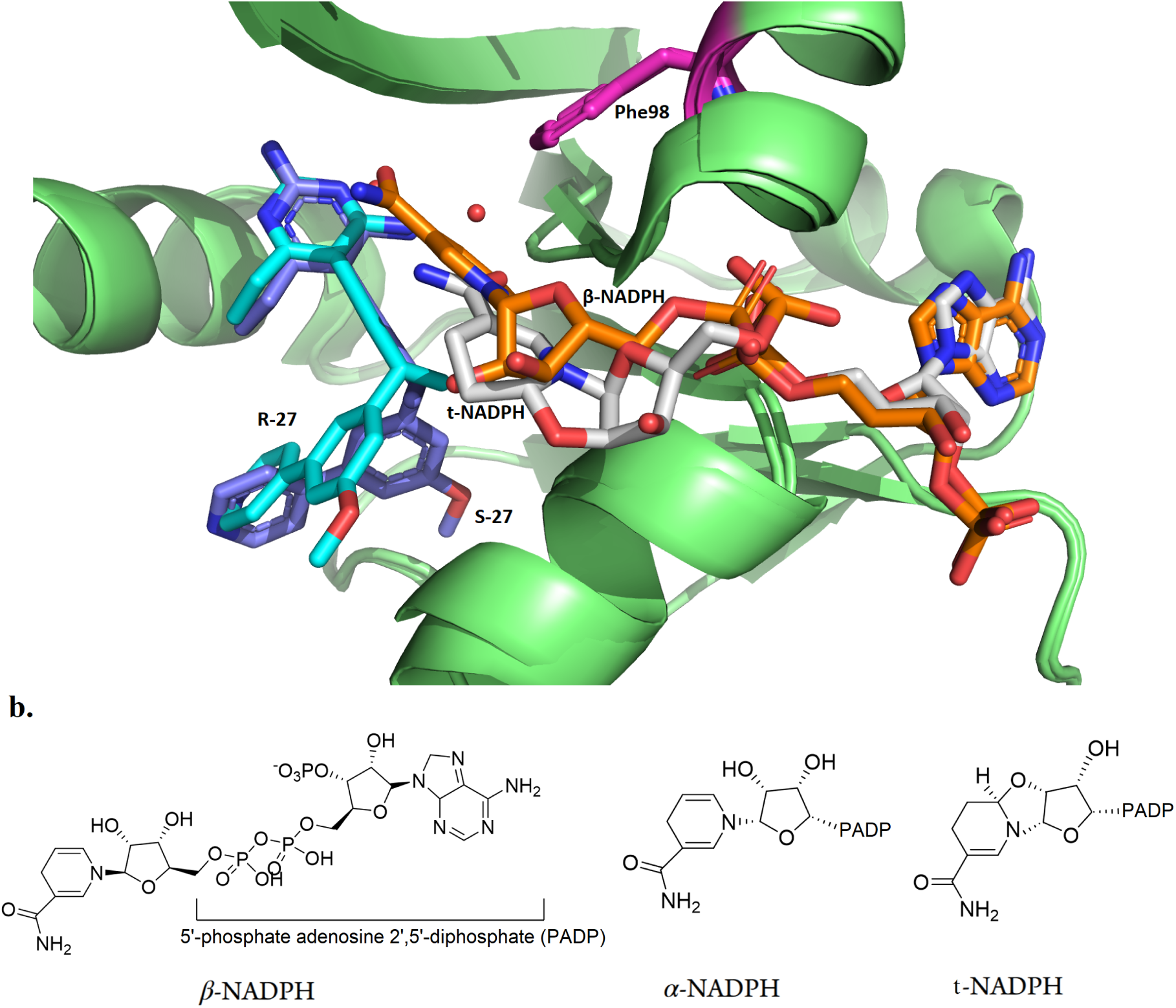
Structure of different configurations of NADPH. **a.** Crystal structures of R-27 (colored in cyan) bound to DHFR and t-NADPH (colored in light grey), and S-27 (purple) bound to DHFR and *β*-NADPH (orange). PDB entries are 6wmy and 4tu5. Phe98 is highlighted in magenta. Starting from the propargylic stereocenter, the biaryl moieties on R-27 and S-27 go to totally different directions and thus R-27 and S-27 adopted very different conformations. **b.** *β*-NADPH, *α*-NADPH and t-NADPH.

In addition to enzymatic analysis, structure determination plays an important role in the study of resistance and development of PLAs. Crystal structures of various PLAs bound to SaDHFR (including both WT and F98Y variants) in complex with NADPH were determined to high resolution (*13*). Interestingly, among all of crystal structures we determined in (*13*), four of them (PDB ID: 3fqf, 3fqo, 3fqv and 3fqz) show two distinct bound forms of the NADPH cofactor. In addition to the common *β*-anomer of NADPH, these structures also contained what appeared to be a second alternative conformation of NADPH. However, subsequently in one of our drug resistance prediction studies for SaDHFR (*17*) using our open-source computational protein design (CPD) software OSPREY (*18–20*), we found that the alternative NADPH model was incorrectly assigned. We thus carefully refitted the density maps of 3fqf, 3fqo, 3fqv, 3fqz using COOT (*21*) and PHENIX (*22*) and performed extensive studies on analyzing the geometric features of the alternative NADPH analog. Over 4000 NADPH conformations binding to different proteins from various species among 1700 deposited PDB files were compared, and we determined that the alternative NADPH ‘conformation’ was in fact a diastereomer with a different anomeric configuration from the standard *β* anomer NADPH (*19, 23–27*). NADPH molecules with different anomeric configurations can naturally interconvert *in vitro* and both forms can be found *in vivo* (*28, 29*).

The stereochemical configuration of molecules is an important factor to be considered in the context of biological processes, particularly in enzyme-catalyzing reactions since enzyme-substrate systems usually require strict chiral matching (*30*). In the SaDHFR system, not only the NADPH cofactor but also the PLAs can bind in different configurations. In order to study how the chirality of the PLA influences inhibitor selectivity and DHFR stereospecifity, a series of PLA enantiomers (determined by a single chiral center on the propargyl linker) were synthesized and analyzed (*16*). In general, most PLA enantiomers pairs can inhibit both WT and F98Y SaDHFR activity at low nanomolar concentration. Interestingly, some PLA enantiomer pairs showed substantially different potencies against SaDHFR. For example, the most potent PLA enantiomer pair R-27 and S-27, have nearly identical IC_50_ values for WT SaDHFR, at 15 and 18 nM respectively. However against the F98Y mutation the R enantiomer suffers a 34-fold loss of potency (IC_50_ 510 nM), whereas the S enantiomer suffers only a 6-fold loss of potency (IC_50_ 111 nM) (Table 1). In other words, S-27 was found to be substantially more resilient towards the F98Y substitution than R-27, despite having almost identical activity towards WT.

To analyze this phenomenon from a structural perspective and to study cofactor influence, two crystal structures of SaDHFR (WT) complexed with R-27/S-27 and NADPH (PDB ID: 4xec and 4tu5) were solved (*16*). According to these crystal structures, we observed that this pair of PLA enantiomers can selectively recruit different configurations of NADPH, which supported our analysis in (*19*). S-27 binds together with *β* form NADPH, while R-27 binds together with another type of NDAPH isomer, with 100% occupancy in both cases (Figure 1a). The alternative NADPH isomer was initially assigned as *α*-NADPH (*16*), but herein we identify it as a tricyclic NAPDH isomer (t-NADPH). NADPH is composed of ribosylnicotinamide 5′-phosphate coupled by a pyrophosphate linkage to the 5′-phosphate adenosine 2′,5′-diphosphate (PADP). The asymmetric center of NADPH discussed here is the 1′ anomeric carbon on the ribose appended to nicotinamide. Three forms of NADPH isomers are shown in Figure 1b. Although these observations suggested a close correlation between resistance and cofactor influence, the detailed mechanism behind it remained unknown.

In this paper, we describe how we first refined the X-ray diffraction data of R-27:t-NADPH:SaDHFR(WT) and updated its structure. Upon careful investigation of the density map, we reassigned *α*-NAPDH in (*16*) as the ring closed, tricyclic tautomer of NADPH (t-NADPH) and deposited the new structure into the PDB (PDB ID 6wmy). Using this new structure, we then applied a series of computational tools and methods to elucidate the mechanism of resistance in F98Y SaDHFR mutants.

The major computational tool we used is our free and open-source CPD software suite OSPREY (*18*). Given a structural model as input, OSPREY exploits provable algorithms to calculate a K* score, which approximates the binding affinity of the given complex (*18*). OSPREY also outputs an ensemble of predicted lowest energy conformations of the protein-ligand complex (*17,31,32*), which provides a detailed model to understand the structural basis of binding affinity changes. In this technique, the thermodynamic ensemble is efficiently predicted by OSPREY, and used to compute K* score which is an approximation of association constant K_*a*_. The output molecular ensemble provides a structural model and explanation for the observed experimental data and computational predictions. Herein, computational techniques allowed us to enumerate and fully explore all possible complexes between different PLA enantiomers, NADPH isomers and SaDHFRs (WT vs. F98Y mutant). Excitingly, our computational results showed excellent concordance with IC_50_ data. The K* scores and ensembles of SaDHFR:cofactor:inhibitor structures calculated by OSPREY suggest an explanation for the stereospecific and cofactor-dependent inhibition of SaDHFR. R-27 and S-27 are an enantiomeric pair of molecules, but when they bind to DHFR they have different preferences for the selection of NADPH isomers. This suggests that there is some form of cooperativity of binding between R-27, S-27 and NADPH, which may be similar to what is described for TMP and NADPH (*33, 34*). Moreover, since R-27 and S-27 have differential potency to F98Y SaDHFR (Table 1), we name the phenomenon *chiral evasion*, which occurs when an enzyme exploits the configuration and chirality difference of its cofactor to evade an inhibitor. In most commonly-seen drug resistant systems, resistance-conferring mutations in enzymes ablate binding of inhibitors and this mechanism is independent of chirality (e.g., TMP resistence in F98Y DHFR doesn’t involve with chirality, since TMP is not a chiral molecule). However, in the chiral evasion case against PLA enantiomers we present in this paper, chirality played an important role in the mechanism of action. There is manifest stereospecific cooperativity between NADPH cofactors and PLA enantiomers, which is supported by crystallographic structure data and the computational results. The OSPREY-produced ensembles of conformations predicted the detailed molecular contacts around the enzyme active sites, providing a structural basis for the mechanism of F98Y-mediated resistance and SaDHFR’s chiral evasion.

The following contributions are made in this work:

1. We reanalyzed the crystallography data of R-27:NADPH:SaDHFR complex, and found that an alternative isomer of the NADPH cofactor namely the tricyclic NADPH (t-NADPH) is a better fit to the original data. Therefore, we present a new structure deposition for this new model (PDB ID: 6wmy). It provides our analysis (below) with an accurate structural foundation.
2. We perform a set of computational analyses to study the PLA enantiomers’ potency to inhibit SaDHFR.
3. K* scores calculated by OSPREY recapitulate the IC_50_ data and explained the reason for PLA enantiomers’ NADPH configuration preference.
4. We propose a mechanism of F98Y-mediated resistance and SaDHFR’s chiral evasion, supported by predicted conformational ensembles.

## 2 Result and discussion

### 2.1 Reassignment of the crystal structure of t-NADPH in complex with DHFR and R-27

NADPH has been observed in different binding states while in complex with SaDHFR and antifolates. In our early study of PLAs in (*13*), an alternative configuration of NADPH that is different from the standard form (*β*-NADPH) was initially assigned to several crystal structures of WT and F98Y SaDHFR (PDB ID: 3fqf, 3fqo, 3fqv and 3fqz). It was hypothesized that this alternative NADPH configuration could be involved in the drug resistance mechanism of SaDHFR (*13*). At that time the alternative form of NADPH was thought to be merely a conformational difference, albeit a unique one. Because of its importance, we then decided to investigate and model this non standard NADPH retrospectively using our CPD software OSPREY. The OSPREY-based analysis (*19*) revealed that the alternative NADPH reported in (*13*) actually has a different configuration from standard *β* - NADPH at the anomeric center. Upon further investigation (*19*), we carefully examined the density map of 3fqf, 3fqo, 3fqv, 3fqz, and measured the geometry (i.e., bond angles) of alternative NADPH around the chiral center (C1^1^ of ribose on nicotinamide side). We searched for similar geometry among over 1700 PDB structures, but only very few were found and among them none was identified that had high enough resolution to conclude anything regarding their anomeric configuration. We also compared to the literature (*35–38*) reports of an NMR study of an alternative NADPH complexed with *Lactobacillus Casei* DHFR (which has the same fold as SaDHFR). Unfortunately, we still found that this alternative NADPH conformation reported in *L. Casei* DHFR study (*35–38*) is distinct from the alternative NADPH we found in our SaDHFR study (*13*). In summary, we found that the alternative NADPH in 3fqf, 3fqo, 3fqv, 3fqz in (*13*) is in a diastereomeric state, and this analog of NADPH had never been reported to bind with non-SaDHFR or non-PLA inhibitors (*19*).

In a later study (*16*) we determined crystal structures of SaDHFR (WT) and NADPH bound to R-27 and S-27, respectively (PDB ID: 4xec and 4tu5). In particular, R-27 was found to bind to this same alternative NADPH configuration. It was indeed confirmed to be in a different isomeric state from *β*-NADPH, which validated our analysis in (*19*). At that time, the NADPH in R-27 complex was thought to be in its *α* configuration (*α*-NADPH, as shown in Figure 1b). However, in this paper, upon review and refinement of the crystal structures we now reveal that the NADPH density is actually in a *α*-02′-6B-cyclotetrahydronicotinamide adenine dinucleotide configuration (t-NADPH), as shown in Figure 1b and Figure 2.

**Figure 2:**
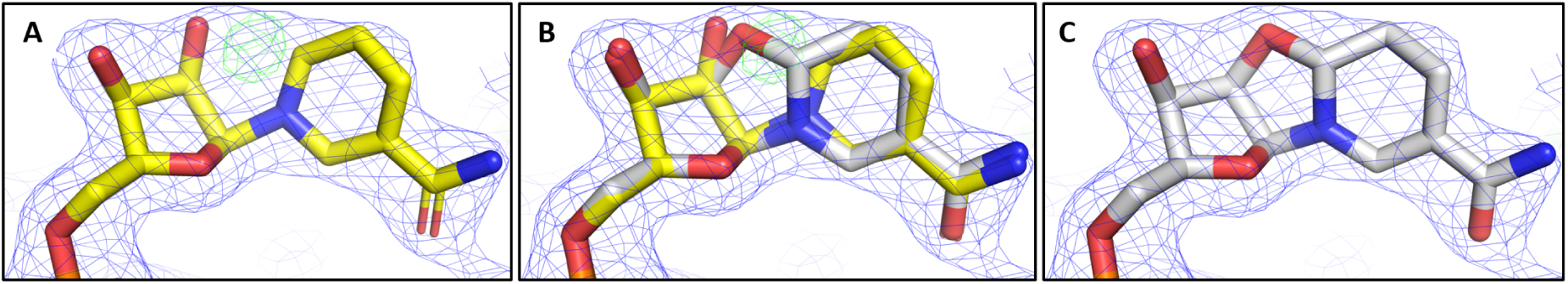
NADPH anomers in the electron densities. **a.** Refinement of *α*-NADPH (yellow) in the SaDHFR:R-27 crystal structure. **b.** Overlay of *α*-NADPH and t-NADPH (light gray) in the SaDHFR:R-27 crystal structure. **c.** Refinement of t - NADPH in the SaDHFR:R-27 crystal structure.

In order to assess the t-NADPH fit in the SaDHFR:R-27 crystal structure, both *α*-NADPH and t-NADPH were refined once into to the final 4xec structure against the original electron density data to eliminate any bias from previous refinement cycles. Upon refinement with the t-NADPH, we saw a reduction in the R_*free*_ from 0.2543 with *α*-NADPH to 0.2529 with t-NADPH (as shown in Table 2). This reduction in R_*free*_ indicates that t-NADPH better satisfies the experimental data and supports the presence of t-NADPH in the active site. Both structures were deposited in the PDB for comparison. The PDB ID for R-27:*α*-NADPH:SaDHFR is 6wmx and the PDB ID for R-27:t-NADPH:SaDHFR is 6wmy. Crystallographic data collection and refinement statistics are provided in Table 2.

**Table 2:** Crystallographic structure data collection and refinement statistics.

| | SaDHFR:R-27:α-NADPH | SaDHFR:R-27:t-NADPH |
| --- | --- | --- |
| PDB ID | 6WMX | 6WMY |
| Space group | P6<sub>1</sub>22 | P6<sub>1</sub>22 |
| No. monomers in asymmetric unit | 1 | 1 |
| Unit cell (a, b, c in Å) | 78.88, 78.88, 107.9<br>90, 90, 120 | 78.88, 78.88, 107.9<br>90, 90, 120 |
| Resolution (Å) | 28.86-2.20 | 28.86-2.20 |
| Completeness % (last shell %) | 97.9 (84.0) | 97.9 (84.0) |
| Unique reflections | 10393 | 10393 |
| $\langle I/\sigma \rangle$ (last shell) | 80.81 (1.62) | 80.81 (1.62) |
| $R_{work}/R_{free}$ | 0.1841/0.2543 | 0.1840/0.2529 |
| No. of atoms (protein, ligands, solvent) | 1406 (1280, 80, 46) | 1406 (1280, 80, 46) |
| Rms deviation bond lengths (Å), angles (deg) | 0.008, 0.89 | 0.008, 0.89 |
| Average B factor for protein (Å<sup>2</sup>) | 37.06 | 37.10 |
| Average B factor for ligand (Å<sup>2</sup>) | 36.22 | 35.99 |
| Average B factor for solvent molecules (Å<sup>2</sup>) | 37.68 | 37.66 |
| Residues in most favored regions (%) | 97.42 | 97.42 |
| Residues in additional allowed regions (%) | 1.94 | 1.94 |
| Residues in disallowed regions (%) | 0.65 | 0.65 |
| Collection location | BNL NSLS X25 | BNL NSLS X25 |

*α* and *β* NADPH are both found in cells *in vivo*, with the *α* form being present at roughly 1.5% of the concentration of the *β* form (*28, 29*). They naturally interconvert although the *β* form is more stable and DHFR active (*28, 29*). However, at low pH *α*-NADPH can become trapped as t-NADPH, by undergoing a cyclisation through the reaction of the 2^1^ hydroxyl on the ribose and tetrahydronicotinamide ring (*39, 40*).

One explanation for the presence of t-NADPH is the low pH (pH 6) used during crystallization conditions. Alternatively, the enzyme binding pocket could potentially alter the pKa of NADPH, thus allowing for the cyclization reaction to occur. The biological relevance of t-NADPH, or its very presence in cells is not known so far. However, to date, t-NADPH has only been found with SaDHFR bound to certain antifolates, and more commonly in SaDHFR mutants containing the resistance mutation F98Y. There are other structures of NADPH complexed with DHFR of different speices in the PDB that were crystallized at a similarly low pH (PH ≤ 6), e.g. 2fzi (*41*), 3frd (*42*), 3fy8 (*43*) etc., but no instances of t-NADPH had ever been found. This suggests that the binding of t-NADPH may play a role in the resistance to antifolates conferred by the F98Y substitution (*16*). In Section 2.3 and Section 3 of the paper we use our comprehensive suite of biochemical, structural and computational data to examine the role of t-NADPH.

### 2.2 Comparison between K* scores and IC50 values

In our previous study (*16*) we found that SaDHFR, in particular its F98Y mutant, showed remarkable stereospecifity of inhibition (Table 1). IC_50_ values of a series of PLA enantiomer pairs were determined, and crystal structures of the most potent pair S-27 and R-27 complexed with SaDHFR and cofactor NADPH were determined. The crystal structure revealed an interesting phenomenon that S-27 and R-27 bound preferentially with different isomeric states of NADPH. S-27 binds with the most commonly seen form *β*-NADPH while R-27 binds with a very rare form t-NADPH. Although the above observations were made, the mechanism of F98Y mediated resistance, the reason of its stereospecifity to enantiomer pairs and the reason for the enantiomers’ selectivity to NADPH configurations remained unclear.

To investigate and understand the above questions, we performed computational analysis using our CPD software suite OSPREY. In order to fully cover all the possible situations, we ran OSPREY on 8 different instances, generated by combinatorially selecting one inhibitor from S-27 or R-27, one NADPH configuration from *β*-NADPH or t-NADPH, one enzyme from WT or F98Y DHFR. The generated 8 instances are listed in Table 3. The source of input starting structures and setup parameters for OSPREY are described in Section 5.2. The resulting K* scores calculated by OSPREY are shown in Table 3. In these cases, the bound state refers to the ternary complex of DHFR:NADPH:inhibitor. The binary complex of DHFR:NADPH as a whole is regarded as unbound state protein receptor, and the inhibitor by itself is unbound state ligand (as explained in Section 5.1 and 5.2).

**Table 3:**
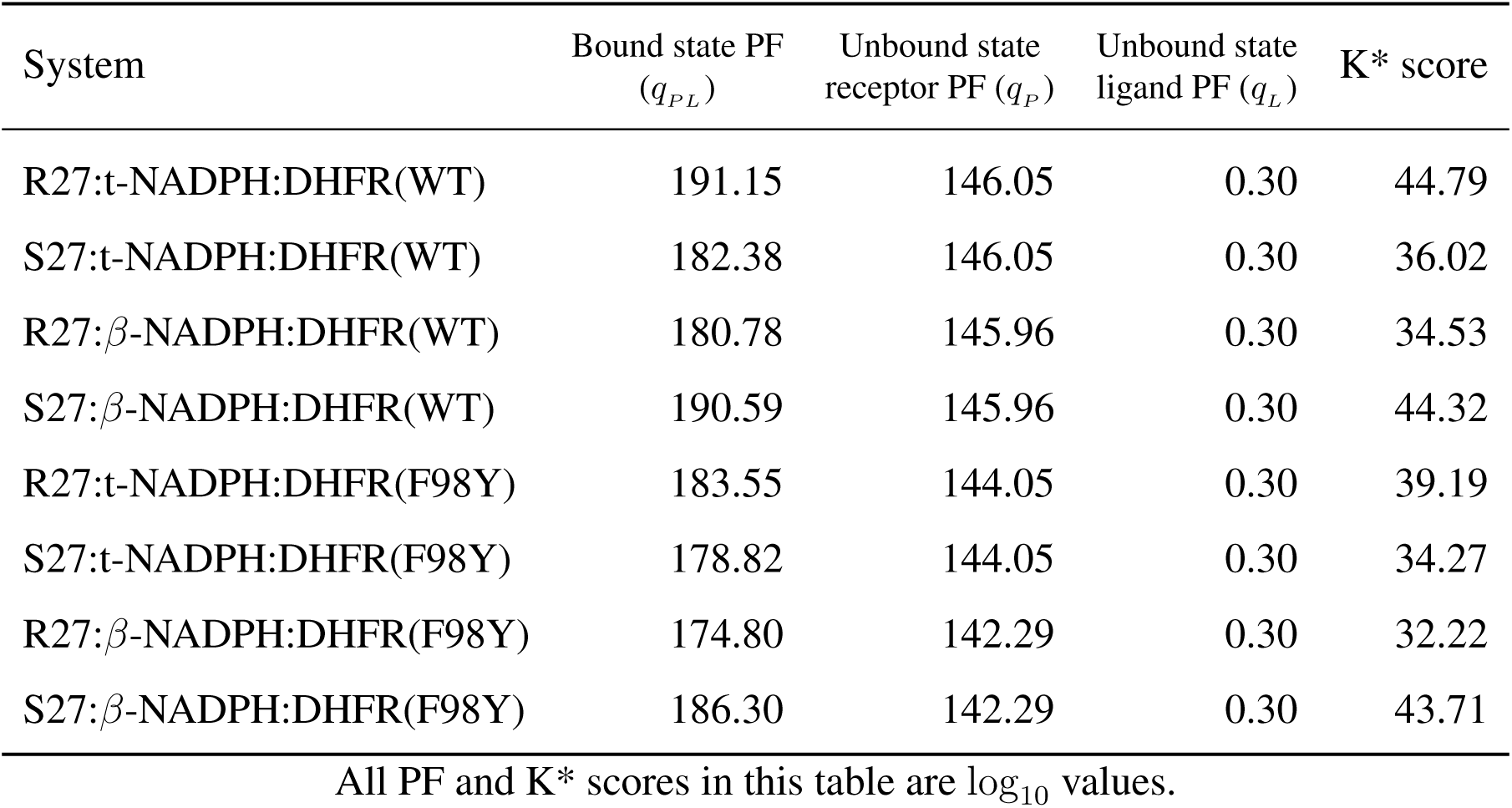
Values of K* scores and partition functions (PF) for different systems calculated by OSPREY.

| System | Bound state PF<br>($q_{PL}$) | Unbound state<br>receptor PF ($q_P$) | Unbound state<br>ligand PF ($q_L$) | $K^*$ score |
| --- | --- | --- | --- | --- |
| R27:t-NADPH:DHFR(WT) | 191.15 | 146.05 | 0.30 | 44.79 |
| S27:t-NADPH:DHFR(WT) | 182.38 | 146.05 | 0.30 | 36.02 |
| R27:β-NADPH:DHFR(WT) | 180.78 | 145.96 | 0.30 | 34.53 |
| S27:β-NADPH:DHFR(WT) | 190.59 | 145.96 | 0.30 | 44.32 |
| R27:t-NADPH:DHFR(F98Y) | 183.55 | 144.05 | 0.30 | 39.19 |
| S27:t-NADPH:DHFR(F98Y) | 178.82 | 144.05 | 0.30 | 34.27 |
| R27:β-NADPH:DHFR(F98Y) | 174.80 | 142.29 | 0.30 | 32.22 |
| S27:β-NADPH:DHFR(F98Y) | 186.30 | 142.29 | 0.30 | 43.71 |

All PF and $K^*$ scores in this table are $\log_{10}$ values.

K* score is an approximation of the binding constant K_*a*_. It is the quotient of the bound and unbound state partition function (PF), where PF is a summation of Boltzmann weighted energy value of all conformations in the design ensemble. Let *q* represent PF, *PL* represent bound state, *P* and *L* represent unbound states protein and ligand, then the definition of K* can be written as 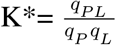 (more detailed description is in Section 5.1). K_*a*_ is an reciprocal of the inhibition constant K_*i*_, and we can show that it is very comparable to IC_50_ in this case. The Cheng-Prusoff equation describes the relationship between K_*i*_ and IC_50_ (*44*). The equation for competitive inhibition case (equation 3 in (*44*)) is:

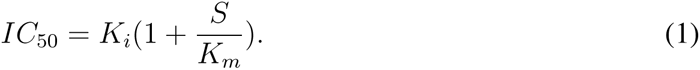

Here K_*m*_ is the Michaelis constant of the substrate and S denotes substrate concentration. For a given type of enzyme and substrate, the value of S and K_*m*_ remain constant, thus K_*a*_ is proportional to 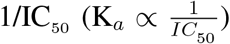. According to the previous study, K_*m*_ values for WT and F98Y DHFR is measured as 14.5 *μ*m and 7.3 *μ*m respectively, and S is approximately 100 *μ*m (*45*). With these data we can convert IC_50_ of our systems into K_*a*_, and then compare to K* scores. The result is shown in Figure 3.

**Figure 3:**
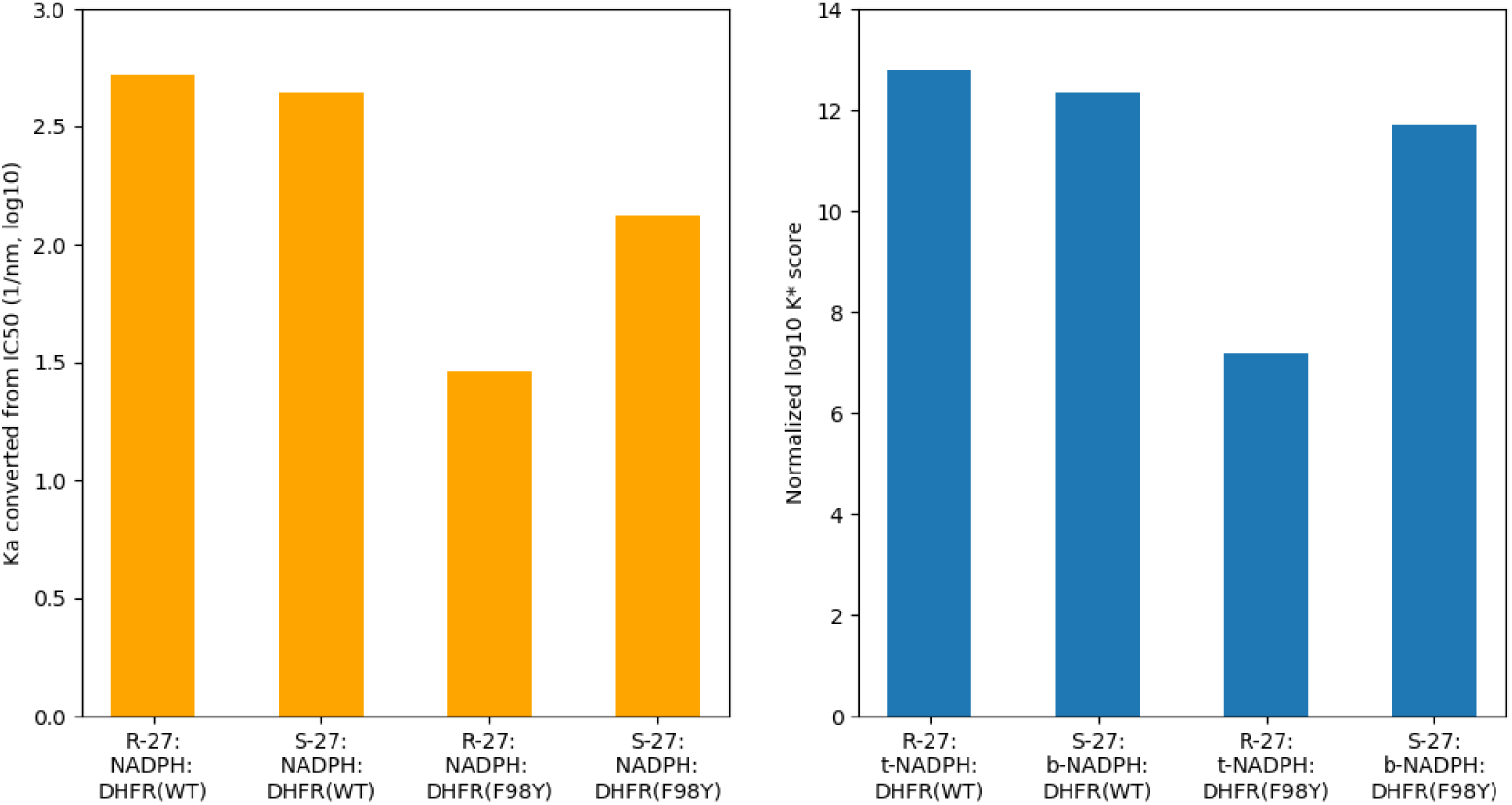
Comparison of K_*a*_ (derived from IC_50_ data) and normalized log_10_ K* score. Bar graphs of K_*a*_ and K* score values (both in log_10_) for different systems. K* scores are normalized as described in Section 5.3. Normalized log_10_ K* scores are in good concordance with K_*a*_, which is derived from IC_50_ using Cheng-Prusoff equation.

From Table 3, we can see that for WT SaDHFR, K* score for R-27:t-NADPH:DHFR(WT) is almost equal to but very slightly higher than for S-27:*β*-NADPH:DHFR(WT). They are much higher than K* scores for S-27:t-NADPH:DHFR(WT) and R-27:*β*-NADPH:DHFR(WT). Such result agrees with IC_50_ data very well. It also indicates that R-27 complexed with t-NADPH is more energetically favorable than when complexed with *β*-NADPH, and on the contrary S-27 is more energetically favorable when complexed with *β*-NADPH than when complexed with t-NADPH. This can explain the difference in NADPH isomeric state observed in crystal structures.

For SaDHFR F98Y, although no crystal structure bound with R-27 or S-27 has been determined, we made homology models (as described in Section 5.2) to investigate their properties. Based on our models, OSPREY predicts that R-27 and S-27 demonstrate the different NADPH configuration preference, as they did when bound with WT SaDHFR. Similarly to those with WT DHFR, K* scores for R-27:t-NADPH:DHFR(F98Y) and S-27:*β*-NADPH:DHFR(F98Y) are significantly higher than those for S-27:t-NADPH:DHFR(F98Y) and R-27:*β*-NADPH:DHFR(F98Y). Moreover, K* scores for S-27 bound with F98Y DHFR only decreased to a moderate extent, compared to when binding with WT DHFR (the log_10_ K* score for S-27:*β*-NADPH:DHFR(WT) is 44.32 and for S-27:*β*-NADPH:DHFR(F98Y) is 43.71, decrease in log_10_ K* score between WT and F98Y is 0.61). But for R-27, compared with bound with WT DHFR, K* score decreased significantly when binding with F98Y (the log_10_ K* score for R-27:t-NADPH:DHFR(WT) is 44.79 and for R-27:t-NADPH:DHFR(F98Y) is 39.19, decrease in log_10_ K* score between WT and F98Y is 5.60). To determine the concordance of K* scores with IC_50_ data, we normalized the K* scores as described in Section 5.3. These normalized K* scores showed remarkably good concordance with IC_50_ data (see Figure 3). Based on these results, our computational analysis successfully recapitulated the experimental thermodynamic and structural data.

### 2.3 Structural analysis

OSPREY can predict and calculate not only the K* score for each protein-ligand complex, but also an ensemble that consists of a series of conformations ranked in the order of energy. These ensembles and conformations exhibit very detailed contact information around the active site, and examining these structures helps us understand the structural basis of F98Y resistance as well as the stereospecific inhibition of DHFR.

Lowest energy conformations from ensembles of 4 wild type SaDHFR-related complexes (R-27:t-NADPH:DHFR(WT), R-27:*β*-NADPH:DHFR(WT), S-27:*β*-NADPH:DHFR(WT) and S-27:t-NADPH:DHFR(WT)) are shown in Figure 4, respectively. The interaction between antifolates (R-27 or S-27) and DHFR and NADPH is visualized using Probe dots (*46*). Red and yellow dots represent unfavorable overlap, and green and blue dots represent H-bonds and van der Waals (vdW) contacts. In all of our analyses, two water molecules in the binding pocket were modeled along with t-NADPH in all complexes containing t-NADPH, since these water moelcules are crucial for bridging contact between t-NADPH, antifolates and DHFR.

**Figure 4:**
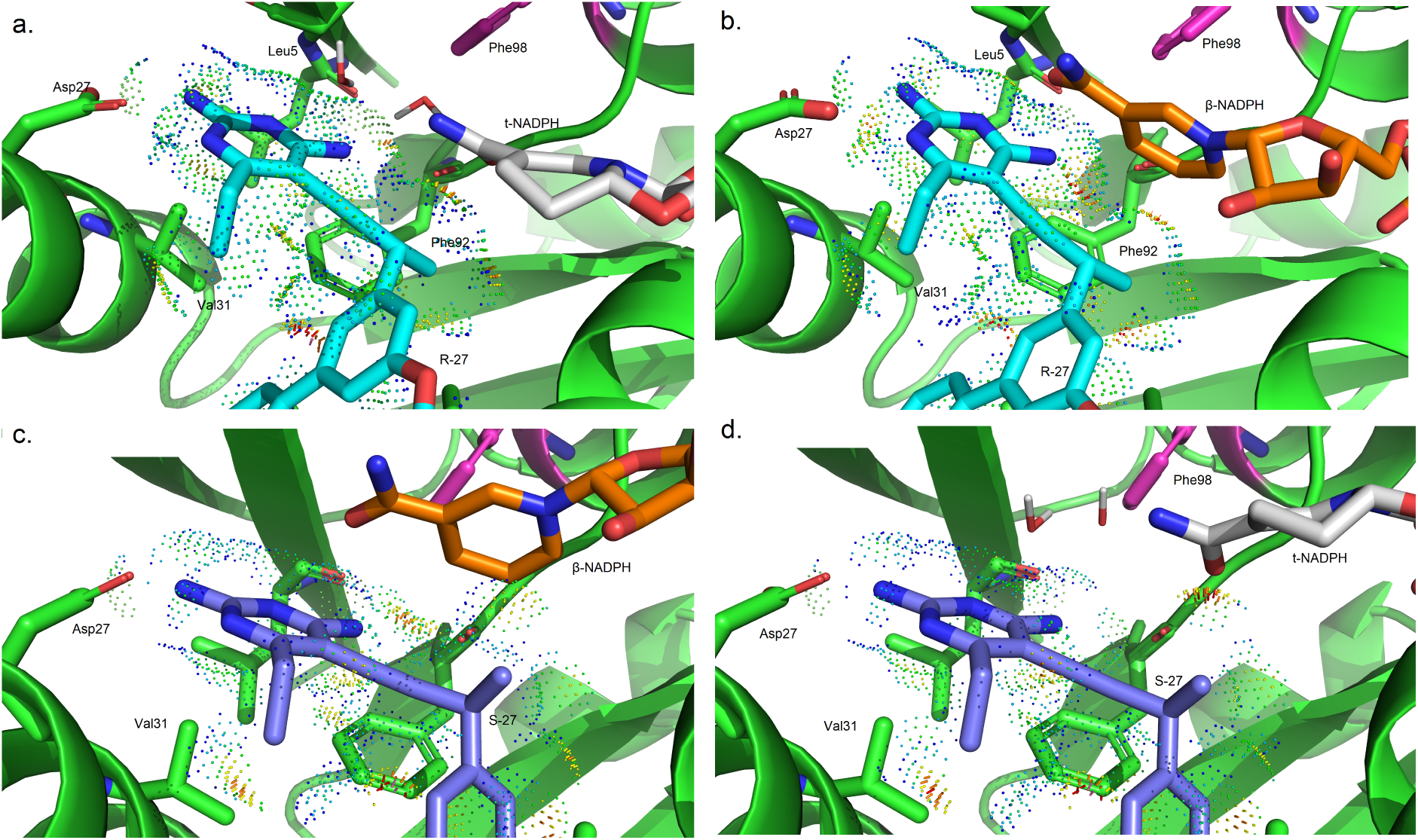
OSPREY-predicted low energy conformations for ternary complexes. **a.** R27:t-NADPH:DHFR(WT) **b.** R27:*β*-NADPH:DHFR(WT) **c.** S27:*β*-NADPH:DHFR(WT) **d.** S27:t-NADPH:DHFR(WT)

As shown in Figure 4, in R-27:t-NADPH:DHFR(WT) system, there are many favorable interactions between R-27 and t-NADPH (panel a). Replacing t-NADPH with *β*-NADPH (panel b) produces some steric clashes between *β*-NADPH and R-27, since *β*-NADPH extends deeper in the binding pocket relative to t-NADPH. It suggests that R-27 binding with *β*-NADPH is less thermodynamically stable than binding with t-NADPH, thus we saw a higher K* score for R-27 binding to t-NADPH and they showed a 100% occupancy in the crystal structure. For S-27, there are more favorable interactions between S-27 and *β*-NADPH (panel c). However, when t-NAPDH is modeled in the same location (panel d), it diminishes many of these favorable contacts. Therefore, S-27 is more stable with *β*-NADPH, and this could explain why S-27 prefers a different NADPH configuration relative to R-27.

To better understand these stereochemical preferences, we performed a structural comparison between the WT and F98Y DHFR binding to R-27 (together with t-NADPH) and S-27 (together with *β*-NADPH) (see Figure 5 and 6, respectively). In R-27 related complexes (Figure 5), compared with the WT DHFR (panel a), the F98Y mutation introduces steric clashes between Tyr98 and Gly93 (panel b). Since tyrosine is bulkier than phenylalanine in size, its side chain is hard to fit in the very confined space around the active site. Clashes induced by F98Y mutation mainly happened between the side chain of Tyr98 and the backbone of Gly93, making it energetically unfavorable. Here the closet distance between Tyr98 and Gly93 is 2.3 Å, which is much closer than what is expected for ideal van der Waals contacts. Panel c and d in Figure 5 showed the lowest energy conformation in binary complex ensembles of t-NADPH:DHFR(WT) and t-NADPH:DHFR(F98Y). According to these binary complex conformations, when no inhibitor binds, significant space in the pocket is left unoccupied. The two water molecules mediating molecular contact for t-NADPH then tend to move outwards to fill up the empty space. Consequently, clashes between Tyr98 and Gly93 that resulted from the crowded space in the ternary complex can be relieved to a large extent in the binary complex. The distance between Tyr98 and Gly93 increases from 2.3 A° (Figure 5b) to 2.6 A° (Figure 5d). The energy and binding affinity change resulted from such difference can be reflected by PF and K* scores. As we can see in Table 3, the bound state PF (log_10_) value of R-27:t-NADPH:DHFR(F98Y) is much lower than R-27:t-NADPH:DHFR(WT) (183.55 vs. 191.15 in log_10_ value), but their unbound state PF value of is quite close (144.05 and 146.05 in log_10_ value). Therefore, their K* scores (as the quotient of bound and unbound state PF) differ (39.19 vs. 44.79 in log_10_ K* value).

**Figure 5:**
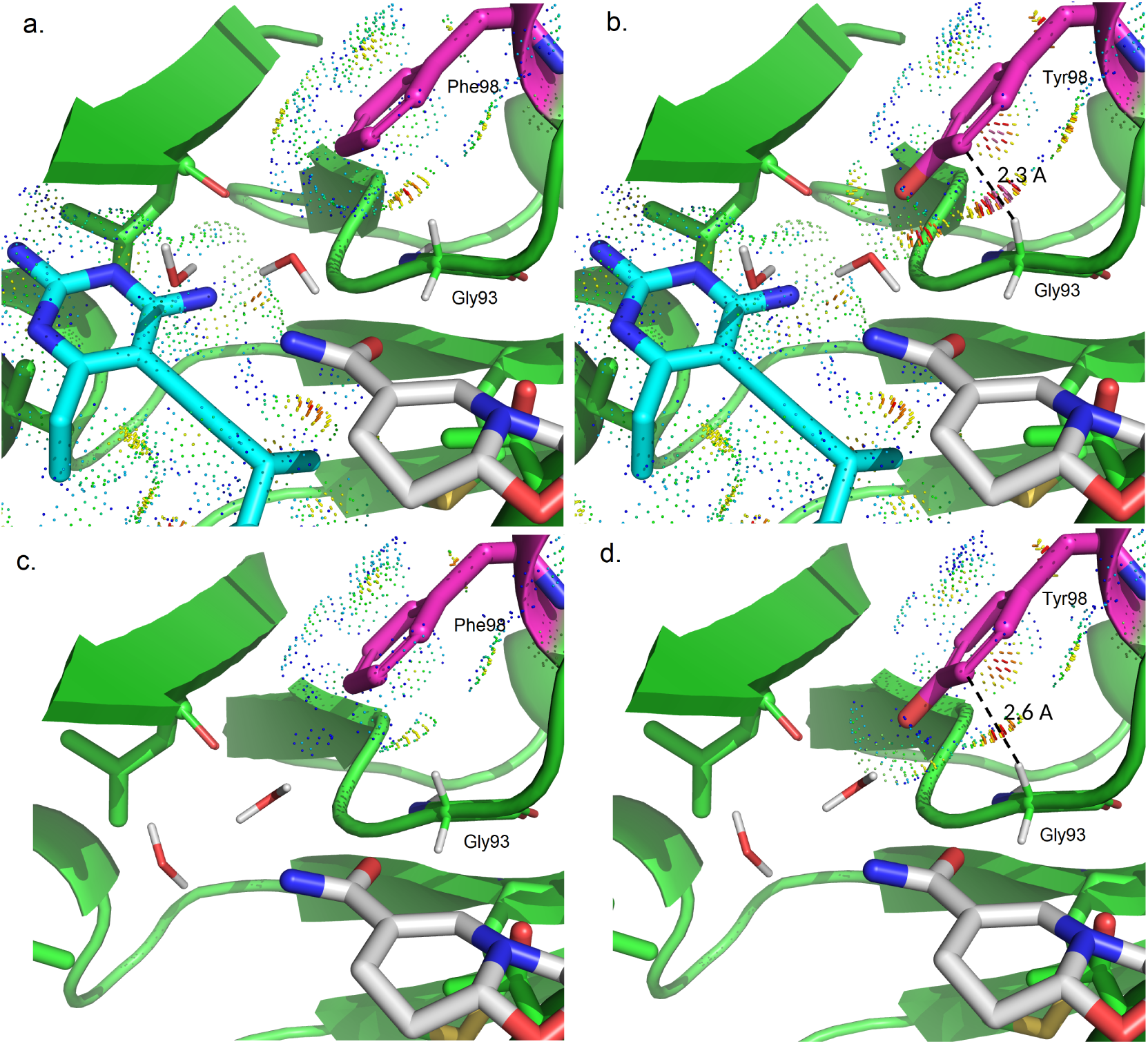
OSPREY-predicted low energy conformations for t-NADPH related complexes. **a.** Ternary complex of R27:t-NADPH:DHFR(WT) **b.** Ternary complex of R27:t-NADPH:DHFR(F98Y) **c.** Binary complex of t-NADPH:DHFR(WT) **d.** Binary complex of t-NADPH:DHFR(F98Y)

**Figure 6:**
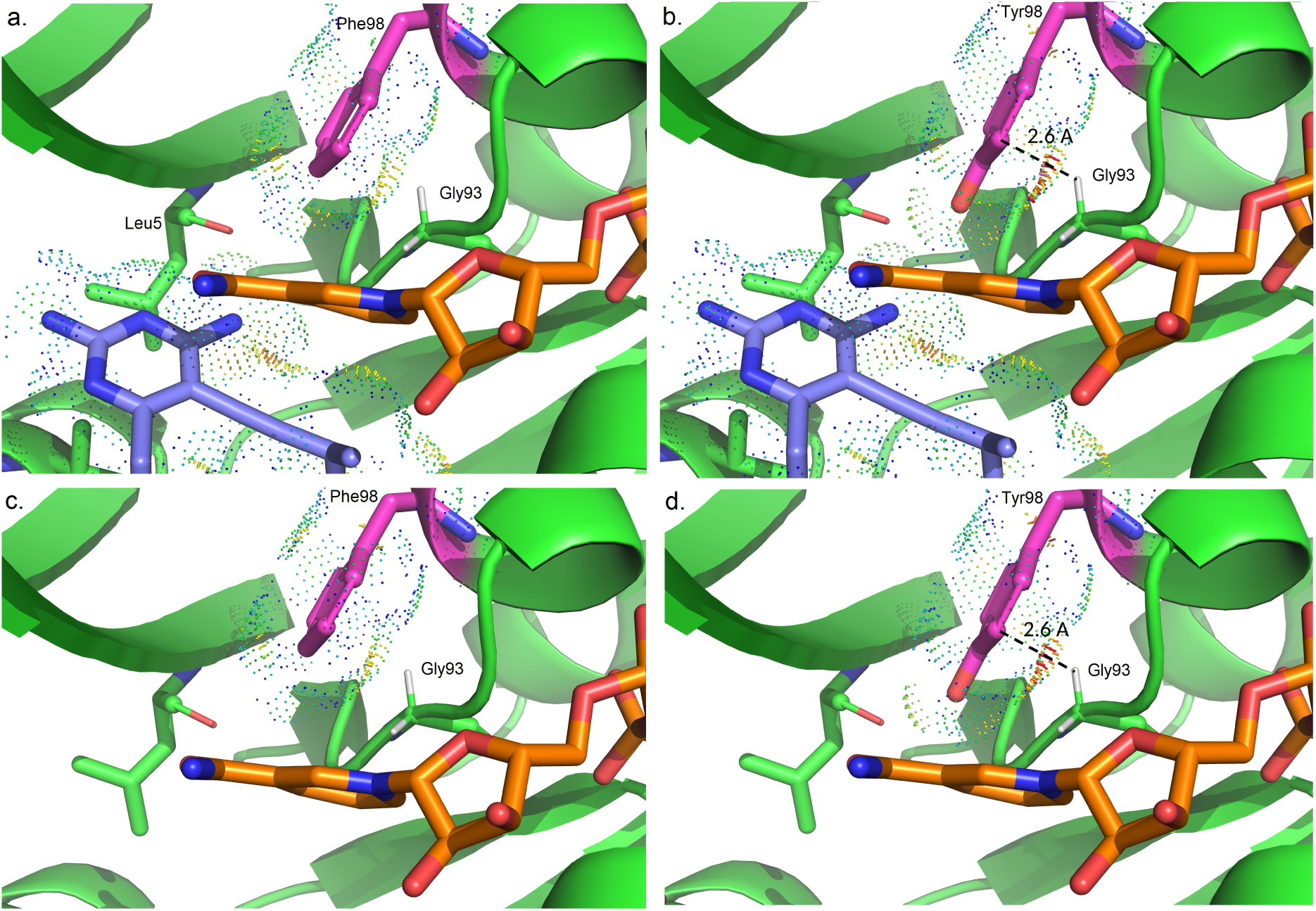
OSPREY-predicted low energy conformations for *β*-NADPH complexes. **a.** Ternary complex of S27:*β*-NADPH:DHFR(WT) **b.** Ternary complex of S27:*β*-NADPH:DHFR(F98Y) **c.** Binary complex of *β*-NADPH:DHFR(WT) **d.** Binary complex of *β*-NADPH:DHFR(F98Y)

For S-27 binding complexes (Figure 6), the influence of the F98Y mutation (panel a and b) is very similar to that with R-27. Compared with the WT amino acid at residue 98 (phenylalanine), the mutation to tyrosine leads to clashes with Gly93. However, the binary complex conformations for *β*-NADPH:DHFR(WT) and *β*-NADPH:DHFR(F98Y) (panel c and d) are very different with those with t-NADPH. *β*-NADPH itself forms substantial contact with DHFR and unlike t-NADPH, there isn’t any water involved in bridging interactions. Even in the binary complex, *β*-NADPH is very restricted such that clashes between Phe98 and Gly93 can not be relieved. Quantitatively, comparing the WT and F98Y DHFR binding to S-27 and *β*-NADPH, PF of both bound and unbound states decrease by similar amounts. Bound state PF values of S27:*β*-NADPH:DHFR(WT) and S27:*β*-NADPH:DHFR(F98Y) are 190.59 and 186.30 respectively, and unbound state PF are 145.96 and 142.29, all in log_10_. Since K* score is the quotient of bound and unbound state PF, when bound and unbound state PF decrease by a similar amount, the decrease in unbound state PF will compensate for the decrease in bound state PF. As a result, (log_10_) K* score did not change too much (44.32 for S27:b-NADPH:DHFR(WT) and 43.71 for S27:b-NADPH:DHFR(F98Y)).

## 3 Discussion

Ability of DHFR to bind different configurations of NADPH could play an important role in SaDHFR’s F98Y-mediated resistance to individual PLA enantiomers, and it could be the key factor that leads to chiral evasion of SaDHFR. Under this model, the ability of DHFR to readily from binary complexes with t-NADPH could be synergistic with the emergence of the F98Y mutation. This is consistent with the observation that the binding of *β*-NADPH to DHFR cooperatively increases the affinity of TMP for DHFR, and that the F98Y mutation disrupts this cooperative binding effect to a considerable degree (*47*). Because of the reorganization of cofactor binding site we showed in this work, TMP would not benefit from cooperativity in this case. In this work we explore our hypothesis by computationally predicting the preference of WT DHFR and the F98Y mutant to two different PLA enantiomers (R-27 and S-27) in the presence of two configurations of NADPH (t-NADPH a nd *β* - NADPH). We specifically use the K* algorithm in OSPREY, a mathematically provable, ensemble-based algorithm that approximates the K_*a*_ value for a complex according to the input model (*18, 31, 32*). Here we show that K* scores recapitulate IC_50_ measurements with a high accuracy (Figure 3). In addition to the complexes observed in previously solved crystal structures, we also made alchemical models that force PLAs to bind together with their unfavorable configurations of NADPH (viz., R-27 with *β*-NADPH and S-27 with t-NADPH). OSPREY predicted the K* scores of these models to be significantly lower than those observed in crystal structures (R-27 with t-NADPH and S-27 with *β*, respectively), indicating these alchemical models are significantly less stable. In this way, we verified that due to the difference of the configuration at the propargyl position, R-27 and S-27 showed different thermodynamic stability when they form complex with different NADPH isomers. Consequently, R-27 and S-27 have a different preference in the NADPH configuration they are binding.

Our *in silico* results provide a putative mechanism for F98Y mediated resistance in SaDHFR, and explain the observed change in IC_50_ of R-27 and S-27. These results indicate that in contrast to S-27, which competes with DHF to bind DHFR:*β*-NADPH, the mechanism of inhibition of R-27 may come from its ability to bind and trap SaDHFR with the inactive t-NADPH. This result is corroborated by the crystal structure of R-27 bound to DHFR (*16*) and our *in silico* results. In both WT or F98Y DHFR, R-27 is predicted to bind to t-NADPH:DHFR with higher affinity than to *β* - NADPH:DHFR (Table 3). Therefore, once R-27 binds to DHFR it will very likely recruit t-NADPH. We believe that t-NADPH is most likely not an active cofactor for three reasons. First, according to our crystal structures (6wmy and 4tu5), it can be seen that the nicotinamide ring of t-NADPH moves farther away from active sites compared to that of *β*-NADPH. Such loss of contacts can make hydride transfer hard to happen. Second, a previous study in (*25*) has shown that *α*-NADPH is inactive with DHFR in almost all species being tested, and t-NADPH is structurally very similar to *α*-NADPH. Third, t-NADPH is less favorable because it will not aromatize upon hydride transfer. Therefore, binding to t-NADPH causes DHFR to be trapped in an inactive state, and reversion to a fully competent state requires the stepwise exchange of the ligand and cofactor. We compared the OSPREY-predicted ensembles and calculated thermodynamic parameters based on these ensembles to reveal the structural basis of chiral evasion of SaDHFR, as well as PLA enantiomers’ different resistance resilience toward the F98Y mutation. For the ternary complex (bound state), the F98Y mutation decreases partition function (PF) values in all instances (Table 3). However, for the binary complex (unbound state) the effect of F98Y is more nuanced. In these cases, our results show that WT SaDHFR binds to t-NADPH with nearly the same log_10_ PF values as *β*-NADPH (146.05 vs. 145.96, only a 0.09 log_10_ difference, see Table 3). While the resistance mutation F98Y causes crowding in the active site when bound to *β*-NADPH, t-NADPH alleviates the crowding in the active site pocket, as described in Section 2.3. This causes the difference in log_10_ PF to increase from 0.09 to 1.67 (the difference between 145.96 and 142.29 is 1.67, see Table 3). The K* scores, in turn, are calculated based on the quotients of PF values, and they strongly agree with experimental measurements (Figure 3). The proposed mechanism above is illustrated in Figure 7.

**Figure 7:**
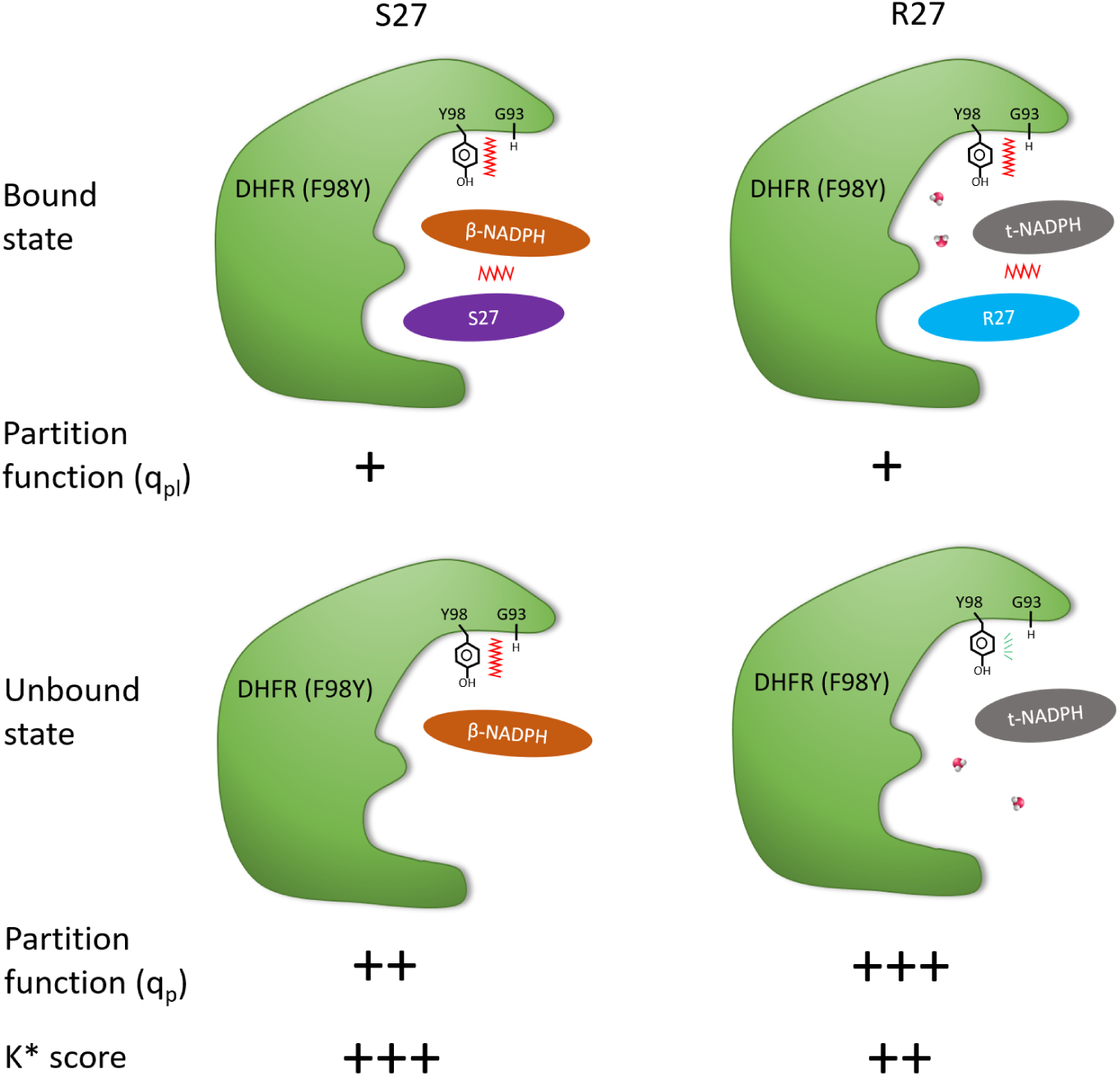
Proposed mechanism for chiral evasion of SaDHFR and resistance resilience by PLAs. R-27 and S-27 have almost equal potency when bound to WT DHFR. However, S-27 binds significantly more tightly to F 98Y DHFR than R-27 d oes. The different configuration of NADPH cofactor played an important role, which we term chiral evasion, according to our data. F98Y mutation introduces steric clashes thus makes the whole complex energetically less favorable. By exploiting chiral inhibitor design, an inhibitor from a PLA enantiomer pair (S-27) was obtained that is resilient to chiral evasion. R-27 binds with t-NADPH and two water molecules, thus the clashes can be relieved in unbound state. In contrast, S-27 binds with *β*-NADPH and the clashes can not be relieved in unbound state. Therefore, as a quotient of bound and unbound state partition function, K* score (which approximates K_*a*_) of S-27 is higher than that of R-27. This suggests S-27 should binds tighter to F98Y DHFR than R-27, which agrees with IC_50_ data. Design for resilience to chiral evasion can therefore overcome drug resistance in a protein target.

In summary, using crystal structure determination, IC_50_ data, and OSPREY prediction we found that in the context of both WT and F98Y SaDHFR, the enantiomeric pair of antifolates R-27 and S-27 bind to different NADPH isomers. We showed this difference in NADPH binding preference leads to the difference in F98Y SaDHFR’s resistance to R-27 and S-27 (S-27 is more resilient to F98Y SaDHFR than R-27). Finally, the structural basis of SaDHFR’s chiral evasion against PLA enantiomers is predicted, as we present in Section 2.3.

## 4 Conclusion

Previous experiments showed that F98Y SaDHFR, a TMP-resistant mutant of SaDHFR, is stereospecific to a pair of enantiomeric PLAs R-27 and S-27. In this study, our CPD software suite OSPREY was employed to explore the mechanism of this stereospecific inhibition, namely, SaDHFR’s chiral evasion against PLA enantiomers. Compared to IC_50_ data, K* scores produced by OSPREY successfully recapitulated the ranking of PLA enantiomers’ affinity and the ranking of the impact the F98Y mutation would have in the interaction. Among all complexes, K* scores (which predict K_*a*_) for R-27:t-NADPH:DHFR and S-27:*β*-NADPH:DHFR are significantly higher than for all other models (Table 3), which is consistent with NADPH configuration preferences observed in crystal structure (R-27 bound with t-NADPH and S-27 bound with *β*-NADPH, as seen in models 6wmy and 4tu5). Ensembles of conformations of F98Y SaDHFR binding to R-27 and S27 were predicted as well. Based on our structural analysis, we found that different binding modes between t-NADPH (which R-27 prefers) and *β*-NADPH (which S-27 prefers) are likely to be a key factor of chiral evasion. The major difference between t-NADPH and *β*-NADPH is how they interact with Gly93 loop on DHFR. Such difference may lead to clashes between Gly93 and Tyr98 in F98Y mutant, and thus may influence the thermodynamic stability of DHFR:NADPH binary complex, ultimately modulating the inhibition potency of R-27 and S-27.

In conclusion we showed how the discovery of a configuration change in NADPH can elucidate a potential mechanism of drug resistance in SaDHFR. This study suggests that the cofactor stereogenicity and chiral evasion should be taken into account when designing new drugs for F98Y SaDHFR. The importance of chirality can not be revealed by merely studying traditional antifolates such as TMP. The use of computational drug design and protein design algorithms gained new insight, informed hypotheses in biology, and made contributions to biochemistry. The data and models we present here have already been useful in medical chemistry campaigns for F98Y resilient inhibitors (*48*), which suggests that they will have significant value for the scientific community.

## 5 Methods

### 5.1 OSPREY and K* algorithm

OSPREY is a structure based CPD software suite developed by us (*18*). For any user-specified mutant sequence of given protein-ligand system, OSPREY is able to calculate a K* score for it which approximates its binding constant K_*a*_. K* score is defined as the quotient of bound and unbound partition function (PF) of the given protein-ligand binding system. The proof of theoretical equality between K* and K_*a*_ under exact accurate condition can be found in Appendix A in (*31*). Bound state refers to the protein:ligand complex, and unbound state is the state when protein and ligand are free and not binding to each other. Let us denote an arbitrary state as *X*, *X* ∈ {*P, L, PL*} where *P*, *L* and *PL* represent unbound state of protein, unbound state of ligand and bound state of protein-ligand complex, respectively. PF of a given state is a summation of Boltzmann weighted energy value for all conformations in the ensemble of this state. For a given mutant sequence *s*, its PF of state *X* (which can be denoted as *q*_*X*_ (*s*)) is defined as:

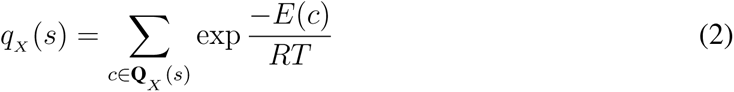

Here **Q**_*X*_(*s*) is the ensemble of state *X*, which consists of every different possible conformation given the sequence *s*. *c* denotes a single conformation **Q**_*X*_(*s*). *E*(*c*) is the conformational energy of *c*. *R* is ideal gas constant and *T* denotes absolute temperature. K* score of *s* the can be written as:

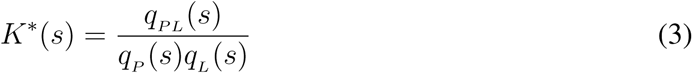

OSPREY uses K* algorithm to calculate K* scores (*31, 32*). The input of K* algorithm includes an starting structure, energy functions, amino acid side chain library and user specified parameters (including flexibility setup, desired mutant sequence, etc.). Under a provable paradigm, it performs A* search (*49*) over the ensemble and generates an ordered, gap-free list of all low energy conformations without any missing. A series of advanced algorithmic techniques, such as *ε*-approximation (a method to bound and approximate PF) and iMinDEE (a method that aims on pruning unfavorable rotamers with continuous flexibility) are also incorporated, making K* algorithm very efficient even handling with large systems (*18*).

### 5.2 Structure preparation and OSPREY setup

OSPREY analysis starts from the input structure, and here are their sources.

In this study, the ternary complex of DHFR:NADPH:PLA is regarded as bound state. For bound state input structures (structures for calculating (*q*_*PL*_)):

- R-27:t-NADPH:SaDHFR: 6wmy
- S-27:t-NADPH:SaDHFR: 6wmy, substituted the original ligand R-27 with S-27
- R-27:*β*-NADPH:SaDHFR: 5ist, subsituted the original ligand (UCP1106) with R-27
- S-27:*β*-NADPH:SaDHFR: 4tu5

Note that 5ist (*50*) was chosen to model the R-27:*β*-NADPH:SaDHFR structure due to the reason that the ligand in 5ist named UCP1106 (also a kind of PLA) adopted a very similar conformation to R-27. Also, for structures containing t-NADPH, 2 key water molecules that bridging contacts near by were included together as well.

For the unbound state structure, since we want to estimate the K_*a*_ of inhibitors, we regard the inhibitor by itself as unbound ligand (for *q*_*L*_) and model the binary complex of DHFR:NADPH as unbound receptor protein (for *q*_*P*_). Input structures for *q*_*P*_ came from:

- t-NADPH:SaDHFR: 6wmy, remove ligand R-27
- *β*-NADPH:SaDHFR: 4tu5, remove ligand S-27

Input structures for *q*_*L*_ are 3D structures of R-27 and S-27 taken from 6wmy and 4tu5, respectively.

All input structures were separately processed through SANDER minimizer from AMBER toolbox (*51, 52*) to release some intrinsic minor clashes in crystal structures. OSPREY is then employed to calculated PF and K* scores. In all of these OSPREY instances, they all have 11 same flexible residues that are Leu5, Val6, Leu20, Asp27, Leu28, Val31, Met42, Thr46, Ile50, Leu54 and Phe92. They have one mutable residue which is Phe98, allowed to mutate to tyrosine, so that we have results for both the WT and F98Y mutant. DHFR, NADPH and inhibitor were modeled with flexibility of translation and rotational movement against each other (*53, 54*). The side chain of flexible and mutable residues were modeled with continuous flexibility (side chain is continuous rotatable within a range of ±9° (*20*) from rotamer library’s discrete value) as well.

### 5.3 Normalization of K* scores

K* scores have been demonstrated many times to be successful in predicting the ranking of K_*a*_ for various mutants of given biological systems (*17,55,56*). However, sometimes the correlation between K* scores and K_*a*_ is not observed to be strictly quantitative (*17, 18*). A few reasons are known to explain such phenomenon: first, our current K* computations only focused on modeling flexibility of resides near the active sites; water molecules out of the binding pocket were not modeled explicitly, but were instead modeled by implicit model (EEF1 model (*57*)); and most physical effective energy functions are based on small-molecule energetics, which can overestimate van der Waals terms. The above limitations in the input model may result in overestimation or underestimation in energy and entropy. In order to compensate for these factors, acquire a better quantitative correlation between K* scores and experimental data and thereby make more accurate predictions, we sometimes normalize K* scores, i.e., scale K* scores of a given system by a constant factor (*56, 58, 59*). In our case here, we can observe from Table 3 that the lowest log_10_ K* score observed among all of our SaDHFR systems is 32.22 (R27:*β*-NADPH:DHFR(F98Y)). Since K_*a*_ must be greater than or equal to zero, we will assume that the zero point of K_*a*_ corresponds to log_10_ K* score approximately equal to 32 here in these cases. Therefore, we calculated a set of normalized log_10_ K* scores, which equals to the original log_10_ K* scores after subtracting an offset value of 32. Note that here we present K* scores in log_10_ space, and a subtraction by constant value in log_10_ space is equivalent to a multiplication in real space.

## Acknowledgements

We gratefully acknowledge grant support from the NIH (R01 GM078031 and R01 GM118543 to B.R.D., and R01 AI111957 to D.L.W.). We thank all the Donald lab and the Wright lab members, Prof. Terrence Oas, Prof. Pei Zhou and Prof. David Richardson for their helpful discussion.

## Author contributions

B.R.D., D.L.W., P.G. and V.G.F. conceived and ideated this project. B.R.D. and D.L.W acquired the funding and provided resources for this study. S.W., A.A.O, M.S.F., G.T.H. and P.G. performed the computational studies. S.M.R. and S.K. provided the experimental data (including the analysis of electron density map and crystal structure determination and deposition). B.R.D. supervised the computational studies and D.L.W. supervised the experimental studies. S.W. and B.R.D. wrote the paper, with comments and contributions from S.M.R., A.A.O., M.S.F., G.T.H., P.G., S.K., V.G.F. and D.L.W.

## Data availability

The crystallography, atomic coordinates, and structure factors have been deposited in the Protein Data Bank, https://www.rcsb.org/ (PDB ID codes 6WMX and 6WMY).

## Code availability

The code of OSPREY (the main computational tool associated with this study) is available at https://github.com/donaldlab/OSPREY3. OSPREY is a free software and is an open source software.

